# Repurposed hnRNPC binds mature mRNAs and safeguards the mitotic transcriptome

**DOI:** 10.1101/2025.09.11.675670

**Authors:** Liat Lev-Ari, Sandra Laster, Andrea Atzmon, Daniel Blumenkrants, Daniel Benhalevy, Orna Elroy-Stein

## Abstract

Upon entry to mitosis, RNA metabolism is broadly suppressed. While mechanistic understanding is limited, the mitotic transcriptome is known to be generally preserved to allow the daughter cells an economical and efficient start. We used highly effective cell cycle synchronization to specifically characterize mitotic hnRNPC, an abundant nuclear RNA-binding protein known for intron binding and splicing regulation during interphase. Two density-distinct hnRNPC-RNP populations (low-and high-density; LD and HD) were identified in mitotic cells. RNA-seq analysis combined with fluorescent Cross Linking and Immuno-Precipitation (fCLIP) revealed hnRNPC binding to 17.1% and 8.7% of expressed genes in LD- and HD-complexes, respectively, with most sites mapping to protein-coding genes (77% and 68%, respectively). Mediated by its known cooperative interaction with U-rich motifs, mitotic hnRNPC acquired prevalent interactions with exons, predominantly within the 3’ untranslated region of mature mRNAs. Mitotic hnRNPC also retained intron interactions, predominantly in LD-hnRNPC RNPs, which comigrated with both the spliceosome and mono-ribosomes through a density gradient. Interestingly, LD-hnRNPC also interacted with mature mRNAs characterized by short coding sequences, in agreement with mono-ribosome loading. Conversely, HD-hnRNPC, which co-migrated with poly-ribosomes, predominantly interacted with mature mRNA complexes. Downregulation of hnRNPC elicited a global negative effect on the abundance of its mitotic targets. The data points to the global role of mitotic hnRNPC as a stabilizer of pre-mRNA and mRNA. Future studies should provide additional insights related to its multifunctional roles during mitosis.

## Introduction

Nuclear envelope breakdown, a fundamental process of cell division, results in a transient loss of subcellular compartmentalization. This event is coupled to the temporary suppression of RNA metabolism, including transcription, splicing (1–5), and translation (6–10), and it is not entirely understood how dividing cells preserve their transcriptome during this phase (9,11–13). Many nuclear RNA-binding proteins (RBPs) participate in splicing during interphase, while their roles following nuclear envelope breakdown remain elusive. Several other RBPs have been shown to participate in distinct biological functions in different cellular contexts (14–23), but comprehensive information about all RBPs is still lacking. Our previous work using quantitative proteomics has shown that mitotic poly-ribosomes associate with a subset of proteins involved in RNA processing, with hnRNPC being the most enriched (24). This observation prompted us to ask what the role of hnRNPC is during mitosis. hnRNPC is an evolutionarily conserved, ubiquitously expressed, and predominantly nuclear RBP first identified as a component of the heterogeneous nuclear ribonucleoprotein (hnRNP) core particle (25). It interacts with intron sequences and accompanies newly transcribed pre-mRNAs during the maturation pathways by the spliceosome (25,26). A previous study applying the <u>C</u>ross-<u>L</u>inking Immuno-<u>P</u>recipitation (CLIP) methodology on non-synchronized cells revealed that hnRNPC, known as a regulator of splicing, binds mostly U-rich sequences within introns, organizing pre-mRNAs into hnRNP particles. This spatially structured binding enables hnRNPC to enhance or repress exon inclusion depending on its position relative to splice sites, highlighting a direct regulatory role for hnRNP particle architecture in splicing control (27). Here, we used a highly effective cell cycle synchronization to specifically characterize mitotic hnRNPC at metaphase. We discovered that mitotic cells contain low- and high-density (LD and HD) hnRNPC-RNPs co-migrating on a sucrose gradient with spliceosomes/monosomes and poly-ribosomes, respectively. Our CLIP data showed that mitoplasm hnRNPC interacts significantly with mature mRNAs, predominantly within HD-hnRNPC-RNPs, recapitulating its molecular preference for cooperative and U-rich binding. Interestingly, LD-hnRNPC also interacted with exons of mature mRNAs characterized by short coding sequences, in agreement with mono-ribosome loading, while it continued to interact with introns associated with spliceosomes. Importantly, we found that downregulation of hnRNPC elicited a global negative effect on the abundance of its mitotic targets. The data points to the global role of mitotic hnRNPC as a stabilizer of pre-mRNA and mRNA.

## Materials and Methods

### Cell culture

Human cervical carcinoma HeLa S3 cells were cultured in Dulbecco’s Modified Eagle Medium (DMEM) supplemented with 10% fetal bovine serum (FBS), 2 mM L-glutamine, and 100 U/mL penicillin (all Biological Industries) at 37°C in 5% CO_2_.

### Cell synchronization to metaphase

Cells were seeded at a density of 8×10^6^ cells per 15 cm plate or 6×10^4^ per well of a 24-well plate. 24 hr later, the cells were treated with final concentration of 2 mM thymidine for 18 hr, followed by three washes with 1x PBS, and addition of fresh medium for incubation of 7.5 hr. proTAME (Cayman chemicals, #25835) was then added to a final concentration of 12 *μ*M for further incubation of 3 hr followed by two washes with 1x-PBS before their harvest for further analysis or fixation for immunofluorescence.

### Sucrose gradient

Cells synchronized to metaphase in a 15cm plate (see above) were incubated with 100 μg/mL cycloheximide (CHX; Sigma, #C7698) for the final 5 min of incubation, followed by ice-cold PBS wash containing 100 μg/mL CHX and, UV-crosslinking of the slightly wet cells using 1500 × 100 μJ/cm² (= 0.15 J/cm²) by placing the plate on ice in a Stratalinker UV Crosslinker (Stratagene). The cells were then lysed on-plate with ‘optimized’ buffer, as detailed in (28), followed by loading onto a 10-50% sucrose gradient for polysome profiling as detailed in (24). 18 fractions were collected along the entire gradient, each containing 0.6 ml

### Immunoblot analysis

Sucrose gradient fractions (see above) were concentrated using StrataClean Resin (Agilent Technologies, cat#400714-61). 10% volume of each fraction was separated by SDS-PAGE followed by Western immunoblot analysis using the following primary antibodies: mouse anti-hnRNPC (clone 4F4, Sigma R5028, 1:1000), mouse anti-PABP (Santa Cruz Biotechnology sc-32318, 1:1000), rabbit anti-RPL26 (Abcam ab59567, 1:4000), rabbit anti-SF2/SRSF1 (Santa Cruz Biotechnology sc-38017, 1:1000), mouse anti-RPS6 (Cell Signaling #2317, 1:1000); and secondary antibodies: HRP-conjugated anti-rabbit IgG, HRP-conjugated anti-mouse IgG, (Jackson ImmunoResearch Laboratories, 1:10 000).

### Immunofluorescence, image acquisition, quantitative analysis

Cells seeded on coverslips (pre-coated with 1 μg/mL PDL for 2 hr at 37°C followed by two 1x PBS washes), in 24-well plates and synchronized to metaphase (see above) were subjected to immunofluorescence staining as detailed in Aviner et al. 2017 (24), using mouse anti-hnRNPC clone 4F4 (Sigma R5028, 1:5000) and Alexa Fluor™ 448 Donkey anti-mouse, (Abcam ab150109, 1:2500) as primary and secondary antibodies, respectively. DNA and actin staining were performed using Hoechst (Sigma #B2261, 1:10,000) and Phalloidin (Thermo Scientific #A12380, 1:250) dyes, respectively. Image acquisition was performed using 3i Marianas (Denver, CO) spinning disk confocal microscope equipped with Yokogawa W1 module, Zeiss Axio-Observer 7 inverted microscope, and Prime 95B sCMOS camera. Objective alpha Plan-Apochromat 100x/1.46 Oil DIC M27 was used, and solid-state diode-pumped 405, 488, and 560 nm lasers; all photometrics under the control of SlideBook™ Intelligent Imaging Innovations. Z-stacks were acquired with a 0.25 μm step size. Images were analyzed using SlideBook 6.0 (Intelligent Imaging Innovations). For each cell, a single midplane optical section was selected to visualize and quantify the spatial distribution and co-localization of the labeled proteins. Only cells in metaphase were included in the analysis based on morphological features. Cell boundaries were manually segmented, and the total cell area (in μm²) was quantified using SlideBook’s area measurement tools. Four subcellular regions were defined within each cell: (i) ‘*DNA region*’, segmented using the Hoechst signal; (ii) ‘*Near DNA*’, defined as a region occupying 20% of the total cell area, surrounding the chromatin mask; (iii) ‘*Periphery*’, defined as a region comprising 33-36% of the total cell area, adjacent to the cell boundary; and (iv) ‘*Remaining*’, defined as the area not included in the previous three compartments. All regional masks were manually adjusted to reflect consistent geometry across cells. Mean intensity for each fluorescence channel was calculated within each region by dividing the total signal intensity by the area of that region. Enrichment was defined as the ratio between the regional mean intensity and the expected mean intensity assuming a uniform distribution across the cell, calculated as the total cellular signal divided by the total cell area: Enrichment_region_=[Mean intensity_region_]/[Total Signal_cell_/Total cell area]. All analyses were performed on a per-cell basis.

### smFISH, image acquisition, quantitative analysis

Cells seeded on coverslips (pre-coated with PDL as mentioned above) in 24-well plates and were enriched for mitosis by a single thymidine block synchronization. After two washes with 1xPBS the cells were fixed by 10 min incubation at room temperature with 4% paraformaldehyde in 1X PBS, followed by two 5 min washes with 1xPBS, and overnight incubation in 70% ethanol at 4°C. Single-molecule RNA detection was performed using DesignReady Stellaris™ RNA FISH Probe Sets ([Biosearch Technologies](https://www.biosearchtech.com/products/rna-fish/designready-stellaris-probe-sets)) designed for RBM3 (Quasar 670, VSMF-2624-5) or POLR2A (Quasar 670, VSMF-2295-5). The smFISH experiments were performed according to the manufacturer’s protocol using the provided buffers, 50nM for each probe, and incubation for 4 hr at 37°C. Coverslips were mounted on slides using ProLong Diamond Antifade Mountant (Invitrogen P36965). Image acquisition as detailed above with the addition of 640nm laser for smFISH spot detection, and analysis using SlideBook 6.0 (Intelligent Imaging Innovations). For each cell, image deconvolution was performed using the nearest-neighbor algorithm across the full z-stack, and a single midplane optical section was selected for further analysis based on maximal cell cross-section. Cell outlines were segmented based on phalloidin staining of cortical actin. Using the phalloidin signal, a binary mask of the total cell was generated. For each cell, the ‘*Periphery*’ compartment was defined based on the phalloidin signal by selecting an inward band corresponding to ∼8–10% of the total measured cell diameter. The width of this region was estimated individually for each cell and adjusted to maintain consistency across the dataset. This manually drawn peripheral region typically represented ∼33–36% of the total cell area. The ‘*Cytosol’* compartment was defined as the remaining cellular area after subtracting the manually defined periphery from the total cell mask. RNA spot detection was performed manually for each cell. An intensity histogram was generated from the RNA channel (e.g., 640nm), and a fluorescence threshold was selected to exclude background signal while retaining discrete RNA spots. This threshold was adjusted per cell to accommodate variation in background intensity and staining efficiency. Spots were segmented using the *Define Objects* function in SlideBook, applying a minimum object size of one pixel to exclude noise. Detected spots were binarized and their xy-coordinates retained. To quantify RNA distribution, each RNA spot mask was tested for spatial overlap with the previously defined ‘*Periphery*’ and ‘*Cytosol*’ masks. Overlapping spots were assigned to either the ‘*Periphery-spots’* or ‘*Cytosol-spots*’ masks, respectively. The number of RNA spots in each region was counted per cell. To account for differences in region size, spot counts were normalized to the relative area fraction of each compartment. Enrichment was calculated as: Enrichment_region_=[%RNA-spots_region_]/[%area_region_]. This enrichment metric was used to quantify the preferential localization of RNA molecules relative to compartment area. All measurements were performed on a per-cell basis.

### Fluorescent Crosslinked-Immunoprecipitation of hnRNPC (fCLIP)

fCLIP, library preparation and initial data analysis were performed as previously described (29) with several modifications. Synchronized HeLa S3 cells were UV-crosslinked on ice (0.15 J/cm²), lysed, and separated by sucrose gradient centrifugation as detailed above. Fractions of interest were pooled (#6-9: Low and #11-17: High-Density fractions). hnRNPC-RNA complexes were immunoprecipitated using a monoclonal anti-hnRNPC antibody (Sigma, R5028) pre-bound to Dynabeads Protein G (Invitrogen), at a ratio of 14.3 μg antibody and 4.3 mg (143 μL) beads per 1 mL of sample. Immunoprecipitation was performed overnight at 4°C, followed by sequential washes in IP buffer (20 mM Tris-HCl pH 7.5, 150 mM NaCl, 2 mM EDTA, 1% NP-40), high-salt buffer (IP buffer containing 500 mM NaCl), and RIPA buffer. Bead-bound RNA-protein complexes were partially digested with RNase I (0.015 U/μL for 10 min at 22°C), then dephosphorylated with QuickCIP (NEB, M0525S), and ligated to a fluorescently barcoded 3′ adapter using T4 RNA ligase 2 truncated K227Q (NEB, M0351). After washing the ligated product was phosphorylated on-beads with T4 PNK, followed by elution with 2X sample buffer and SDS-PAGE. Crosslinked RNA-protein complexes were visualized using a fluorescent imager, and gel slices corresponding to the ligated hnRNPC RNP (∼25 kDa larger than hnRNPC Mw) were excised. 3’-ligated RNA footprints were recovered by proteinase K digestion followed by phenol–chloroform extraction and ethanol precipitation. Recovered RNA was ligated to a 5′ adapter, reverse transcribed using SuperScript IV (Thermo Fisher) and amplified by 5-cycle PCR. Then, following size selection on a 3% agarose Pippin Prep, a second PCR was performed adding indices and the scaffolds required for Illumina NGS. Final libraries were purified (Zymo DNA Clean & Concentrator), quantified by TapeStation, and assessed for adapter-adapter contamination. Libraries were sequenced on an Illumina platform.

Demultiplexed .fastq files (GEO GSE307226) were generated using Bcl2fastq (Illumina) and Cutadapt [--adapter=NNTGACTGTGGAATTCTCGGGTGCCAAGG] (30). hnRNPC binding sites (groups; Supplementary Table S2) were defined by Paralyzer (31) (https://github.com/ohlerlab/PARpipe), and raw data visualization was performed using IGV (32). fCLIP replicates were also derived from alternative experimental conditions (high RNase concentration of 0.15 U/μL) that produced similar results and conclusions (shown in Supplementary Figure S2). For low-versus high-RNase fCLIP analysis, four datasets (HD-Low, HD-High, LD-Low, LD-High) of fCLIP libraries were prepared, processed, and aligned to GRCh38 (Ensembl release 109) as part of Paralyzer pipeline (31). Then, reads per gene matrices were derived by intersecting reads with Ensembl exon annotations using Python/pysam+intervaltree fallback, normalized to counts-per-million (CPM). Genes with CPM ≥0.5 were considered bound. Next, transcripts annotated as *protein_coding* were retained based on Ensembl gene biotype classification (Ensembl release 109(33)). Within-fraction comparisons (low-versus high-RNase) were performed separately for HD and LD fractions, and overlap metrics (Jaccard index, Fisher’s exact test) as well as correlation coefficients (Pearson, Spearman) were calculated. PCA was performed on protein-coding CPM values (prcomp, R), and Low–High concordance within fractions was further assessed by ordinary least squares and robust regression (HuberT) on log1p-transformed CPM values (shown in Supplementary Figure S2(F-J)).

hnRNPC non-targets were defined as expressed genes (RNAseq reads >0) not bound by hnRNPC (CLIP reads=0). Only groups residing in expressed genes with #groups>2 were defined as hnRNPC binding sites (Supplementary Table S3).

Standardized binding intensity by gene: for exons, the 5’UTR, CDS, and 3’UTR reads (see Supplementary Table S3) sum was divided by transcript length; for introns, intron reads per gene were divided by the gene total intron length. Metagene analysis of hnRNPC binding along genomic coordinates was performed using NGSplot (34), with the following parameters: -G hg38 -R exon -L 500 -RB 0.01 -P 0 -FL 34 -SS same -IN 1 -GO km -RR 50 -BOX 0. Metagene analysis of hnRNPC binding along transcriptomic coordinates was performed as in (35). *K-mer* analysis was performed as in (36). Retained Intron analysis used findings from (37) (Table S6, Human introns and properties), including introns of types B and C, and not A), followed by hg19 to hg38 coordinates conversion(38). Intron coordinates .bed files were intersected using bedtool intersect -u(39). High-RNase replicate analysis is shown in Supplementary Figures S2(C-E).

### RNAseq

Prior to loading on Sucrose gradient, samples of the total cell extract were removed (100 µl) for RNA extraction by mixing with 900ul of RiboEX reagent (GeneAll), followed by vortexing, incubation at RT for 5 min, chloroform addition (200 μL) and vortexing, a 2-minute incubation at RT, and centrifugation at 12,000 × g for 15 min at 4 °C. The aqueous phase (450μL) was transferred to a new tube. RNA was precipitated by addition of 1 μL PelletPaint (Merck), 0.1 volumes of 3 M sodium acetate (pH 5.2), and 1.5 volumes of isopropanol, followed by overnight incubation at −20 °C and centrifugation at 20,000 × g for 30 min at 4 °C. Pellets were washed with 1 mL of ice-cold 70% ethanol and centrifuged at 7,500 × g for 5 min at 4 °C. After air-drying for 10 min, RNA was resuspended in 10 μL of 10 mM Tris-HCl (pH 8.0) and incubated at 56 °C for 5 min to ensure complete dissolution. RNA was depleted from rRNA (NEB, E7405) and used for NGS library preparation (NEB, E7760), followed by Illumina sequencing (NextSeq2000, 100 bp, single read).

### Standardization and comparison to RNAseq data from (24)

Raw sequencing reads of DTB-mitosis and DTB-G1 (24) were trimmed using Cutadapt (30) with the following parameters: [--adapter=CTGTAGGCACCATCAATTCGTATGCCGTCTTCTGCTTG, -- minimum-length 20, --max-n 0, --quality-cutoff 20 --cores 2]. Quality of trimmed FASTQ files was assessed using FastQC (40) and alignment to GRCh38 (Ensembl release 109) was performed using STAR aligner (41) (v2.7.11b). Genome index was pre-built using --sjdbOverhang 27, corresponding to read length of 28 bp. Alignment was performed with the following parameters: --runThreadN 8 --outFilterMultimapNmax 20 --alignSJoverhangMin 5 --alignSJDBoverhangMin 1 --outFilterMismatchNmax 2 --outFilterMismatchNoverReadLmax 0.1 -- outFilterScoreMinOverLread 0.33 --outFilterMatchNminOverLread 0.33 --alignIntronMin 20 -- alignIntronMax 1000000 --alignMatesGapMax 1000000 --alignEndsType Local. Gene-level quantification was performed during alignment using STAR’s --quantMode GeneCounts, followed by differential expression analysis using DESeq2 (42). To generate a high-confidence gene set for downstream comparisons, a six-step filtration strategy was applied to the DTB-G1 and DTB-Mitosis RNA-seq datasets, resulting in a final list of 229 protein-coding genes (Supplementary Table S1): (i) Only transcripts annotated as *protein_coding* (Ensembl gene biotype classification, Ensembl release 109(33)) were retained. Genes were then filtered based on DESeq2 differential expression analysis between G1 and Mitosis: (ii) significantly differentially expressed genes (adjusted *p*-value < 0.05) were retained; (iii) genes with stable expression across conditions were retained if they had *padj* > 0.05 and an absolute log_2_ fold change < 0.07. (iv) Genes with no measurable expression in either condition were excluded by removing those with a combined G1 and Mitosis read count ≤1, based on the merged count matrix. (v) To prevent bias inferred by the increased sequencing depth of the metaphase samples, genes with fewer than 400 summed raw counts in the Metaphase dataset were also excluded. This threshold was defined based on the ∼30-fold greater coverage in the metaphase dataset compared to G1 and Mitosis. (vi) Finally, to remove transcripts affected by batch effect between datasets, genes with fold detection ratios (Mitosis/Metaphase) outside the interquartile range were excluded. The final set of 229 genes, selected for consistent detection and absence of major technical artifacts, was used for downstream comparisons and visualization. Although DESeq2 analysis was performed on the full dataset, principal component analysis (PCA) and clustering analysis were restricted to this high-confidence gene set. Gene expression values were variance-stability transformed (VST), and PCA was conducted using the first two principal components. PCA was performed using the prcomp function on VST-transformed, unscaled gene expression values (scale. = FALSE). For heatmap generation, variance-stabilized transformed (VST) expression values were row-centered by gene, and unsupervised hierarchical clustering was performed using the cutreeDynamic function (deepSplit = 1, minClusterSize = 10), resulting in five distinct gene clusters. The full list of clustered genes is provided in Supplementary Table S1.

### Statistical analyses

GraphPad software (Prism version 10.4) was used for plotting and statistical analysis. Figure 1C “*hnRNPC distribution in interphase and metaphase”* was plotted as Box and whiskers (Tukey); P-values were determined by 2-way-ANOVA and Sidak’s multiple comparison test. Figure 1D “*hnRNPC distribution in metaphase 4 regions of the cell”* was plotted as Box and whiskers (Tukey); P-values were determined by 1-way-ANOVA and Sidak’s multiple comparison test. Figure 3C, “*Group intensity by gene”* was plotted as violin plots (truncated); P-values were determined by Kolmogorov-Smirnov test to evaluate either the exon or intron binding intensity of LD vs. HD. Figure 3E “*median CDS length - hnRNPC target vs non-targets LD and HD”* was plotted as median with 95% CI; P-values were determined by Kruskal-Wallis. Figures 4B-D, 6A,B, S4 “*Cumulative Distribution analysis of mRNA length, CDS, 3UTR, 5UTR and hnRNPC KD RPF and RNA levels”* were plotted as Histograms with Nonlin fit; P-values were determined by Kruskal-Wallis.

**Figure 1:**
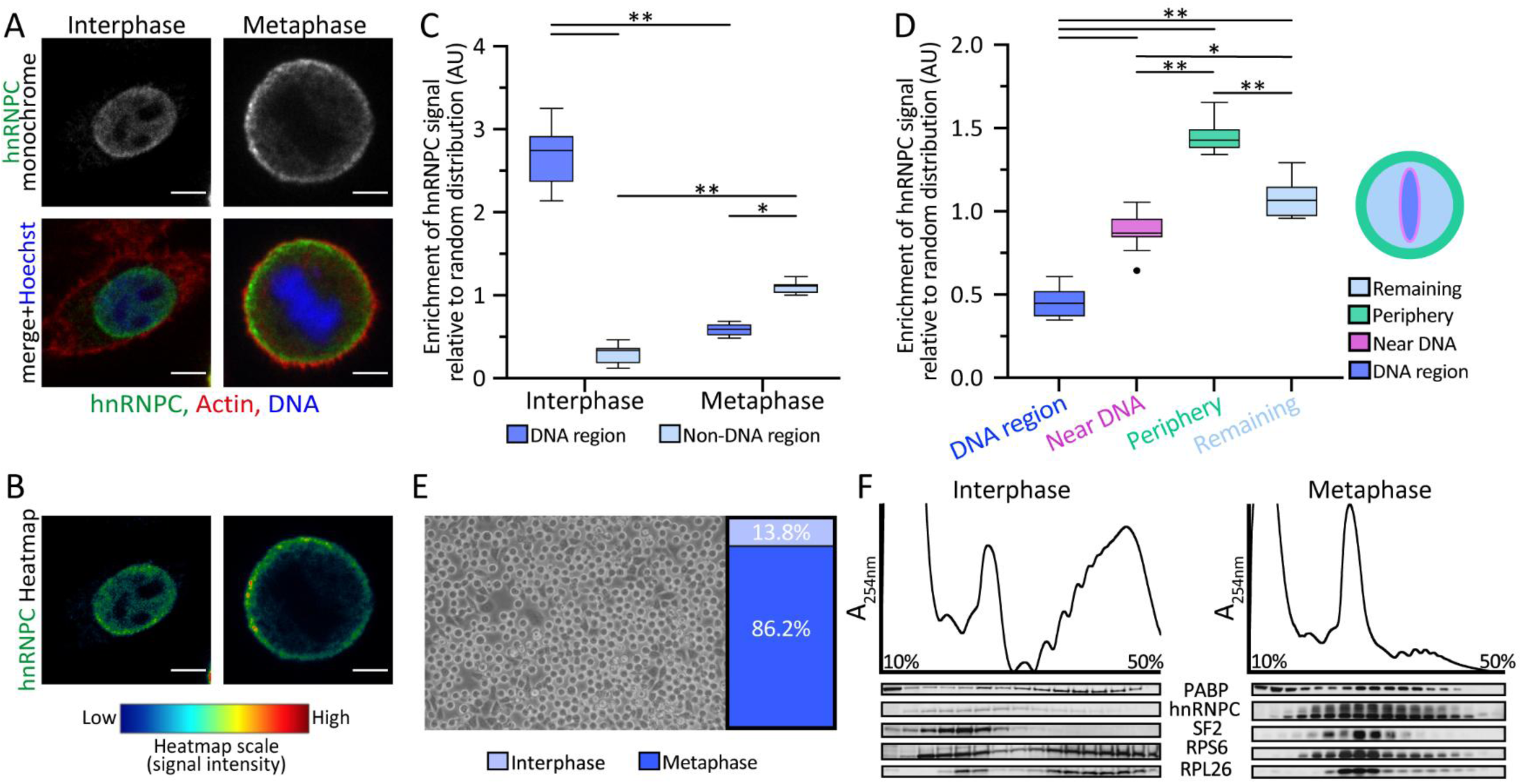
Sub-cellular distribution of hnRNPC at interphase and metaphase. (A) Representative confocal images of HeLa S3 cells in interphase and metaphase stained for actin (phalloidin, red), DNA (Hoechst, blue), and immune-stained for hnRNPC (green). Scale bar, 10 μm. (B) Pseudocolor heatmap showing hnRNPC signal intensity from the same confocal plane as in (A). (C) hnRNPC distribution to DNA and non-DNA regions in cells at interphase (n=12) and metaphase (n=8). DNA regions were segmented based on Hoechst intensity; non-DNA regions were defined by subtraction from the manually segmented cell area. Shown is the enrichment of the hnRNPC signal relative to random distribution. Two-way ANOVA with Tukey’s post hoc test: *P = 0.0002, **P < 0.0001. (D) hnRNPC distribution to four defined sub-cellular regions of cells at metaphase (n=11). One-way ANOVA with Tukey’s: *P = 0.0004, **P < 0.0001. (E) Representative image of HeLa S3 cells after synchronization using a single thymidine block, followed by proTAME treatment. (F) Absorbance (A_254nm_) and Western Blot analysis of non-synchronized (interphase, left) and proTAME-synchronized (metaphase, right) HeLa S3 cells after lysis and 10%-50% sucrose gradient fractionation.

## Results

### Two density-distinct hnRNPC-RNPs are detected during mitosis

To maximize sensitivity and specificity for mitotic hnRNPC interactions, we aimed to work with cells optimally synchronized to a specific time point, when relevant RNPs of interest are most enriched. Since we previously found that hnRNPC predominantly localizes to the cellular cortex in cells synchronized to mitosis by double thymidine block (DTB) (24,43), we assumed this subcellular location reflects functional engagement of mitotic hnRNPC with RNP complexes. Therefore, we utilized this unique subcellular localization phenotype as a readout for mitotic hnRNPC function and determined the time of its full appearance along the mitotic process. We found that while nuclear egress of hnRNPC into the cytoplasm initiates at prophase, its peripheral localization is fully acquired only at metaphase. hnRNPC localization to the cell cortex then persisted through anaphase and telophase, followed by nuclear re-entry during cytokinesis, in concert with nuclear envelope reformation (Figure 1AB and Supplementary Figure S1A). Quantitative analysis of confocal immunofluorescence confirmed hnRNPC co-localization with nuclear chromatin during interphase, and its exclusion from the DNA region at metaphase (Figure 1C). Spatial quantification of four different subcellular areas confirmed hnRNPC-specific enrichment at the cell periphery at metaphase (Figure 1D). To extract RNPs from cells specifically at metaphase, we used proTAME for cell synchronization, given its high effectiveness in synchronizing cells to this specific sub-mitotic timepoint (44). Our optimized synchronization procedure, comprising proTAME treatment following a single thymidine block, consistently attained 85-90% of cells at metaphase (Figure 1E).

As an initial analysis, we wanted to test for differential gene expression in cells synchronized to metaphase by proTAME (current study) relative to cells semi-synchronized by DTB to mitosis or G1, for which RNAseq data was previously obtained, though at lower coverage (24,43). After careful data filtration, we could list 229 genes that were sufficiently detected in all datasets for downstream analyses (see M&M for filtration strategy, Supplementary Figure S1B, C, and Supplementary Table S1). Principal component and clustering analyses of variance stabilized (VST) gene expression (42) clearly demonstrated the higher efficiency of proTAME in cell synchronization relative to DTB, with a mixed-characteristic pattern of gene expression in DTB-synchronized cells (Supplementary Figure S1BC, and Supplementary Table S1). Since DTB synchronization is less selective, it retains a larger portion of non-mitotic cells, while proTAME synchronization is highly effective and yields a more homogeneous cell population and gene expression trend. Genes upregulated in mitosis comprise clusters 1 and 3, and include MDM2 (45), the mitotic spindle associated ZNF263 (46,47) (cluster 1), and genes associated with chromatin organization, a critical process for DNA condensation during mitosis, such as RNF40 (48), HDAC5 (49), and MacroH2A1 (50) (cluster 3). In contrast, cluster 2 consists of genes that are downregulated in mitosis and is enriched with genes related to positive regulation of cell growth and DNA replication, linked to G1 and S phases, respectively (Supplementary Table S1).

We hypothesized that hnRNPC executes its mitosis-related functions while it is associated with specific RNP complexes. Therefore, we employed fractionation of RNPs on a sucrose gradient, as traditionally used for polysome profiling (24), using extracts of non-synchronized and proTAME-synchronized cells to identify mitosis-specific hnRNPC-containing fractions. Immunoblot analysis was applied to the fractions using antibodies specific for SF2 (spliceosomes marker), RPS6, RPL26 (small and large ribosomal subunit markers, respectively), PABP (translation complexes marker), and hnRNPC (Figure 1F). In line with translation downregulation during mitosis (6–10), a smaller polysomal peak was exhibited by cells synchronized to metaphase. As expected from a nuclear protein involved in splicing, hnRNPC was not detected in cytoplasmic fractions in non-synchronized cells; and even cells at metaphase exhibited predominant co-sedimentation of hnRNPC with splicing factor SF2 to sub-polysomal fractions containing the spliceosome complex (51) and mono-ribosomes. Complexes within these fractions were termed low-density (LD)-RNPs. Surprisingly, at metaphase, hnRNPC also migrated to the denser fractions that harbor mRNAs loaded with poly-ribosomes, and do not contain SF2, nor are known to comprise the splicing machinery. Complexes within these fractions were termed high-density (HD)-RNPs (Figure 1F).

### Identification of RNA sequences bound by hnRNPC within LD- and HD-RNPs in metaphase

To identify RNA sequences bound by hnRNPC in metaphase, we applied the fluorescent Cross Linking and Immuno-Precipitation methodology (fCLIP) (29). Cells synchronized to metaphase were exposed to UV (254nm) for protein-RNA crosslinking, followed by cell lysis and sucrose gradient fractionation to isolate the distinct subpopulations of hnRNPC-RNPs. Two samples were generated by pooling the fractions containing either the LD-RNPs (harboring monosomes and spliceosomes) or the HD-RNPs (harboring the polysomes). The LD and HD-RNP pools were each subjected to immunoprecipitation using an antibody specific for hnRNPC and analyzed by fCLIP to capture hnRNPC-RNA interactions, followed by SDS-PAGE and gel-purification of the hnRNPC-RNP of interest (Figure 2; Supplementary Figure S2A). The fCLIP analysis identified 28862 and 12521 binding sites (a.k.a groups (29,52)) in LD- and HD-hnRNPC fractions, respectively. Among these, 77% of LD-hnRNPC and 68% of HD-hnRNPC binding sites mapped to protein-coding genes (Supplementary Figure S2B). The complete lists of binding sites, including those detected along non-coding RNAs (pseudogene, snRNAs, miRNA, lncRNA, and more), are provided in Supplementary Table S2. Integrating fCLIP and RNA-seq analyses revealed that 8.7% and 17.1% of expressed genes are bound by hnRNPC within the HD-RNP and LD-RNP pools, respectively. As a preliminary assessment of the fCLIP procedure, we examined whether LD- and HD-hnRNPC targets recapitulate the well-established preference of hnRNPC for binding U-rich sequences (27,53–55). To this end, we performed k-mer analysis (*k*=6) to identify enriched 6-mer motifs within the hnRNPC-bound sequences. This analysis indicated a clear preference of both LD- and HD-hnRNPC for poly-U tract binding, starting from 4 consecutive Us and peaking at the U 6-mer (UUUUUU) (Figure 3A). This result strongly indicates that authentic hnRNPC interactions were identified by the fCLIP procedure.

**Figure 2:**
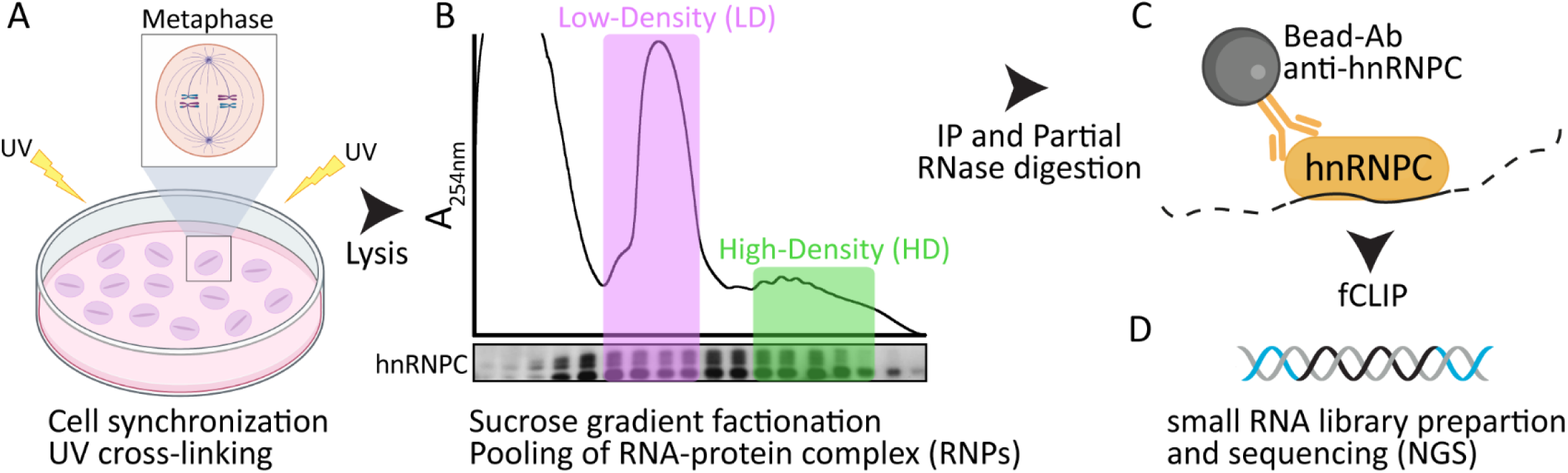
Scheme of experimental setup. (A) Cell synchronization to metaphase followed by UV-mediated protein-RNA crosslinking. (B) Sucrose gradient fractionation and pooling of low-density (LD) and high-density (HD) RNPs. (C) Immunoprecipitation (IP) of LD- and HD- RNP fractions using anti-hnRNPC Ab, followed by partial RNase digestion (dashed line). (D) Completion of the fCLIP procedure, small RNA cDNA library preparation, and next-generation sequencing (NGS).

### Intronic and exonic binding sites are differentially distributed between LD- and HD-RNPs

LD-hnRNPC RNPs co-sediment with the spliceosome marker SF2 to densities previously shown to harbor spliceosomes (51); therefore, it is reasonable to assume that at least part of LD-hnRNPC- containing RNPs are involved in splicing. However, given that the LD fractions contain a mixture of spliceosomes and mono-ribosomes, while mitotic hnRNPC also migrated to denser, poly-ribosomal fractions (HD-hnRNPC) that lack the spliceosome marker SF2, we hypothesized that some of the LD-hnRNPC might interact with mature mRNAs engaged with mono-ribosomes. To test this hypothesis, we assumed that for splicing-related interactions, hnRNPC will target pre-mRNAs, predominantly at introns, as previously shown (27); and for post-splicing -related interactions, hnRNPC will target exons, comprising mature mRNAs. Dissecting the exonic and intronic nature of hnRNPC binding sites showed that, in agreement with its co-migration with SF2, LD-hnRNPC interacts more with introns than HD-hnRNPC (64% versus 44%, respectively). While exonic binding of LD and HD-RNPs is distributed along the entire mRNA, HD-hnRNPC interacts more with 3’UTRs than LD-hnRNPC (38% and 21%, respectively; Figure 3B), indicating that a larger portion of HD-hnRNPC targets were loaded with ribosomes, hindering CDS interactions (56,57). To further compare the functionality and specificity of LD- and HD-hnRNPC targets, we tested whether intronic targets were previously reported as retained introns (37), which therefore may be included in mature mRNAs that co-sedimented with polyribosomes. While only 10% of the introns bound by HD-hnRNPC were identified as retained introns, their frequency was 2.5-fold higher among HD-hnRNPC than LD-hnRNPC-bound introns (Supplementary Figure S3A), thus HD-hnRNPC further exhibited a higher preference for mature mRNAs.

**Figure 3:**
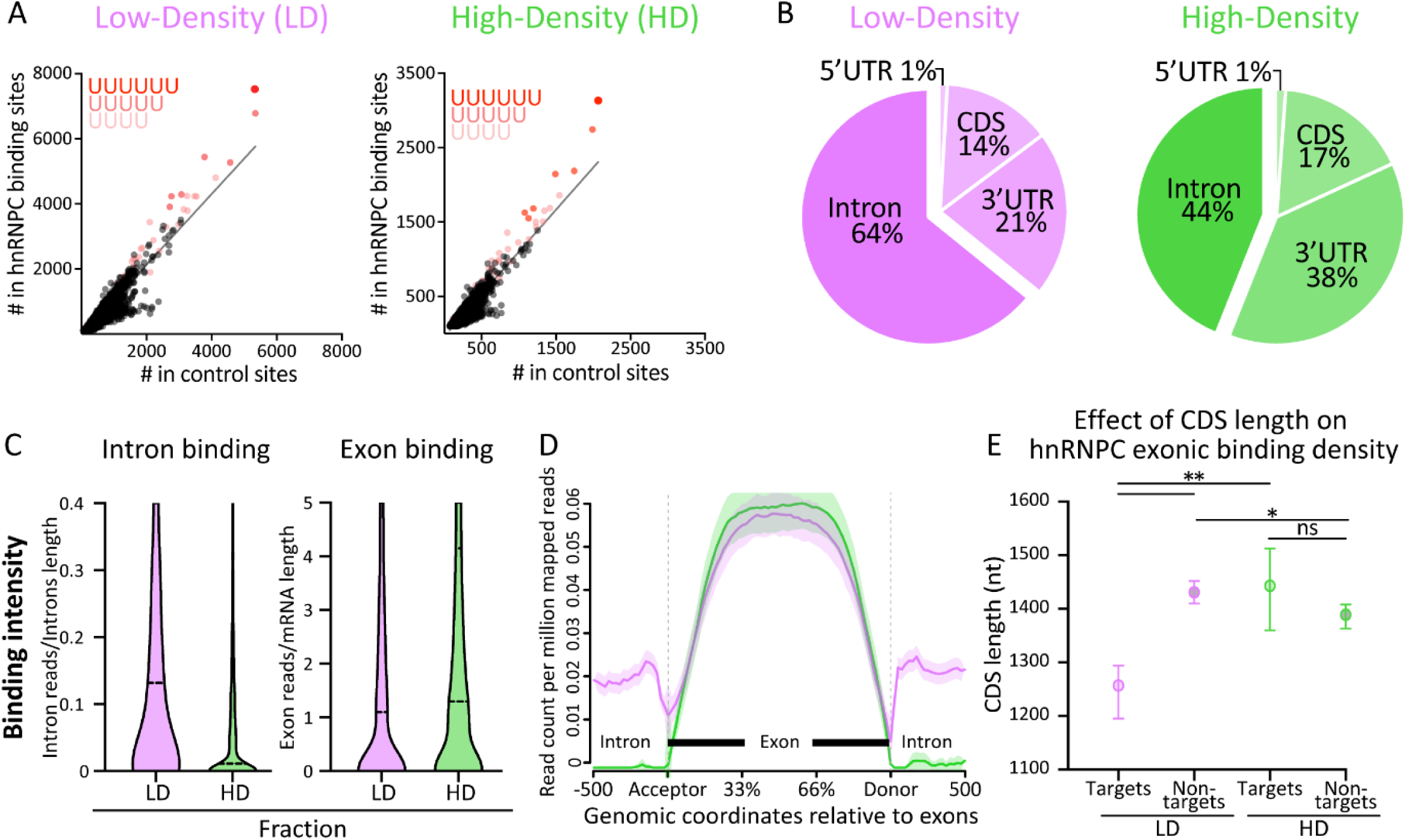
Characterization of hnRNPC binding sites. (A) 6-mer analysis for sequence motif enrichment within hnRNPC-bound regions in LD- and HD-RNP fractions. Scatterplots show observed frequencies of each 6-mer relative to its frequency in a control set. 6-mers containing 4, 5, and 6 consecutive U are color labelled. Information related specifically to exonic and intronic binding is shown in Supplementary Figure S3. (B) Distribution of hnRNPC binding between introns and exons, segmented by mRNA functional elements (CDS-coding sequence, UTR-untranslated region). (C) hnRNPC binding intensity per intronic and exonic binding sites, calculated per gene by normalizing read counts to introns or mRNA lengths. (D) Metagene distribution of hnRNPC binding along exons ±500 nt in the adjacent up- and down-stream introns. HD, green; LD, pink. Shaded areas represent standard error. (E) Coding sequence (CDS) length of hnRNPC exonic targets and non-targets in LD (pink) and HD (green) fractions. Median values ± 95% confidence interval are presented. Kruskal–Wallis test: *P = 0.005, **P < 0.0001; ns, non-significant.

Next, we utilized the quantitative nature of fCLIP to compare the binding intensities of LD- and HD-hnRNPC with introns and exons. As shown in Figure 3C, while LD- and HD-hnRNPC interacted with exons at comparable intensities, LD-hnRNPC interactions with introns exhibited an order of magnitude higher intensity, relative to HD-hnRNPC–intron interactions, suggesting that a large portion of detected HD-hnRNPC intronic binding is at minimal levels and at least in part representing false positives due to fCLIP high sensitivity. Transcriptome-wide positional meta-analysis similarly indicated that only LD-hnRNPC quantitatively interacted with introns (Figure 3D), for which *k*-mer analysis of intronic and exonic targets confirmed that all are characterized by poly-U tract binding (Supplementary Figure S3B). Thus, collectively, our analyses indicate that HD-hnRNPC interacts preferentially with mature mRNAs, while LD-hnRNPC exhibits a more complex pattern of interaction with both exonic and intronic RNA sequences.

LD-hnRNPC mixed interaction with exons and introns and its co-migration with spliceosomes and mono-ribosomes (Figure 1F) raises the question whether LD-hnRNPC complexes comprise a mixture of spliceosomal pre-mRNAs and mRNAs encoding relatively small proteins, whose short coding sequence (CDS) often enables loading of only a single ribosome. To clarify this insight, we compared the CDS lengths of LD-hnRNPC targets to those of HD-hnRNPC targets and expressed non-hnRNPC targets. Notably, only LD-hnRNPC exonic targets were characterized by a selectively shorter length of CDS (Figure 3E). This finding is consistent with LD-hnRNPC comprising a mixed RNP composition of mono-ribosomes loaded with mature mRNAs (58), and spliceosomal pre-mRNA complexes, unlike the HD-hnRNPC RNP composition, which predominantly includes poly-ribosomes loaded with mature mRNA complexes. Given that translational states are affected by mitotic suppression (6,10), hnRNPC association with mono- and poly-ribosomal mRNAs brings up the possibility that these mature transcripts may either be actively translated or engaged with stalled ribosomes.

### fCLIP confirms the minimal-length binding property of hnRNPC

Based on the literature, hnRNPC-RNA interaction is highly cooperative, with single hnRNPC tetramers serving as the core binding unit, and tri-tetramers forming the 19S ‘triangular complex’, binding an RNA stretch of ∼700 nt (59,60) (see Figure 4A, adapted from Figure 12 of Huang et al., 1994 (60)). The ∼700 nt complex formation is considered the functional *in vivo* form of hnRNPC; however, since to-date, hnRNPC has been shown to interact mostly with pre-mRNAs, this assumption could not be tested in cells due to the considerably higher than ∼700 nt intrinsic length of introns. The finding of mitotic hnRNPC interaction with mature mRNAs now provides an opportunity to test the length-dependent cooperative hnRNPC-RNA interaction model. To this end, we focused exclusively on mitotic hnRNPC exonic targets, representing mature mRNAs, and tested for length dependence. As shown (Figure 4B), while hnRNPC non-targets were of lengths down to 350 nt, hnRNPC-bound mRNAs were selectively above 620 nt for LD-hnRNPC and 670 nt for HD-hnRNPC. These findings are in striking agreement with the previous findings, supporting the notion that hnRNPC-RNA interaction is highly cooperative also during mitosis, with tri-tetramers forming the 19S ‘triangular complex’.

**Figure 4:**
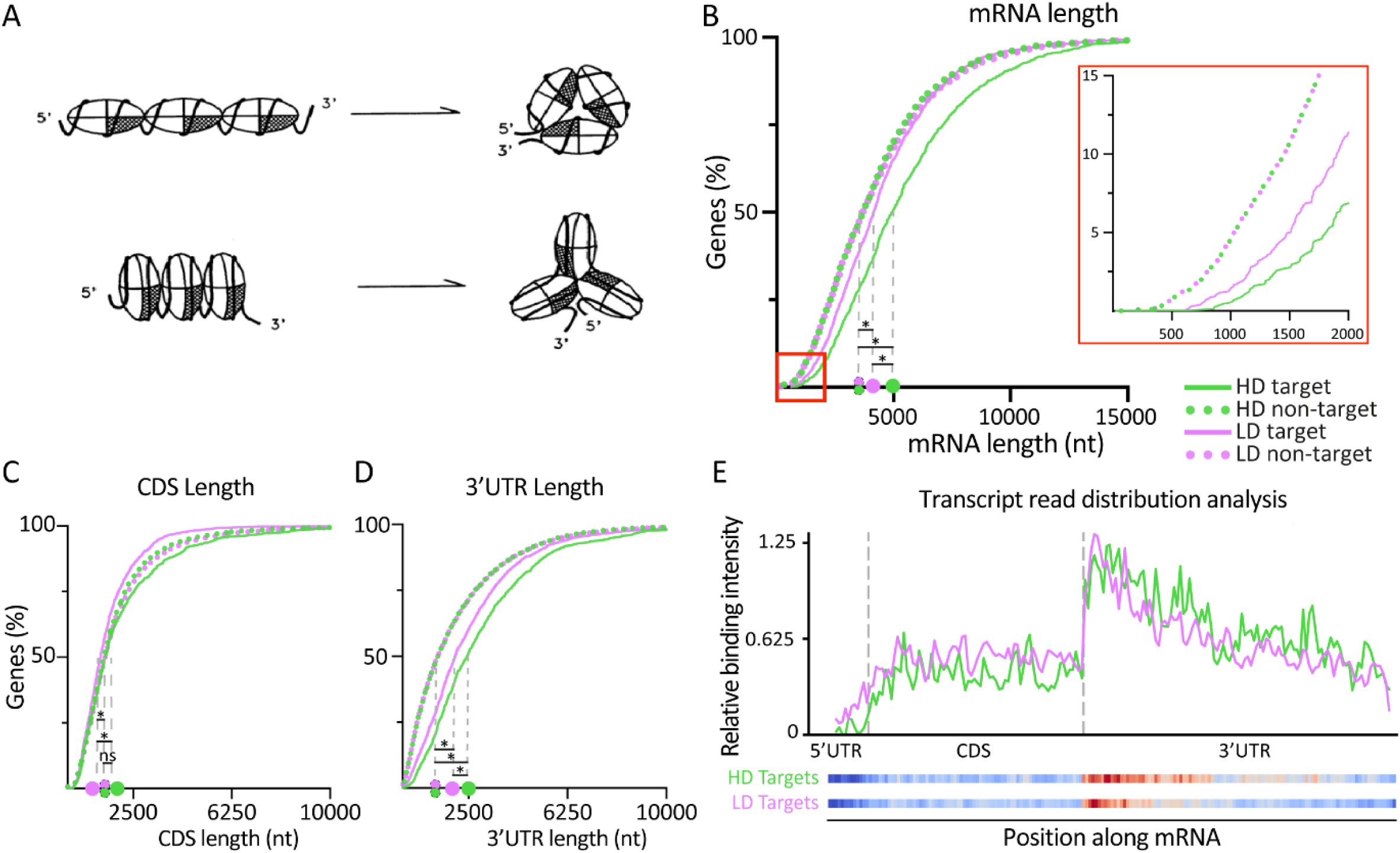
Analysis of hnRNPC length-dependent Exonic Binding. (A) Cooperative hnRNPC binding model, adapted from Figure 12 of Huang et al., 1994 (60). (B–D) Cumulative length distribution of mRNA (B), CDS (C), and 3’UTR (D). HD, green; LD, pink; Targets, continuous line; non-targets, dashed line. Median values are presented on the x-axis. Kruskal–Wallis test: *P < 0.0001; ns, non-significant. (E) Metagene analysis of read distribution relative to mature mRNA coordinates, segmented by 5′UTR, CDS, and 3′UTR.

### hnRNPC-mRNAs interactions exhibit a preference for the 3’UTR

To probe the functionality of mitotic hnRNPC-RNA interactions, we leveraged the apparent minimal-length binding property of hnRNPC. We therefore asked whether, in exon-targets, is it the CDS, 5’UTR, or 3’UTR lengths that limit hnRNPC association. Our analyses indicated that the length of the 3’UTR, but not that of the CDS and 5’UTR, is relevant for exonic binding of hnRNPC, and thus for its interaction with mature mRNA (Figure 4CD, and Supplementary Figure S4). In accordance, meta-transcriptomic positional analysis demonstrates a clear preference for hnRNPC interaction with the 3’UTR (Figure 4E), as would be expected by the higher occurrence of U-rich binding sites within many 3’UTRs (61). This pattern is characteristic of cytoplasmic RBPs-mRNA interactions, which are often more abundant along the 3’UTR, while CDS regions are loaded with ribosomes and less available for RBPs to interact (56,62). Given that hnRNPC has until now been mainly associated with splicing, we aimed to explore the functional significance of hnRNPC interaction with mature mRNA.

### hnRNPC functions to protect the mitotic transcriptome from RNA degradation

Since hnRNPC acquires cell cortex localization during metaphase, we first wanted to corroborate a link between this unique sub-cellular localization and mitotic hnRNPC interaction with mature mRNAs. To this end, we selected two candidate mRNAs for smFISH analysis that are expressed at similar levels: RBM3, an exon-exclusive target of hnRNPC, and POLR2A, an hnRNPC non-target (see IGV visualization of RBM3 and POLR2A in Supplementary Figure S5). As shown in Figure 5AB, RBM3 mRNA was enriched at the cell periphery, in correlation with mitotic-hnRNPC localization, while POLR2A mRNA was depleted from the cell periphery (Figure 5AB; Figure 1A). This result strongly suggests that at least part of the mitosis-acquired hnRNPC interactions and putative functions are taking place at the cell cortex.

**Figure 5:**
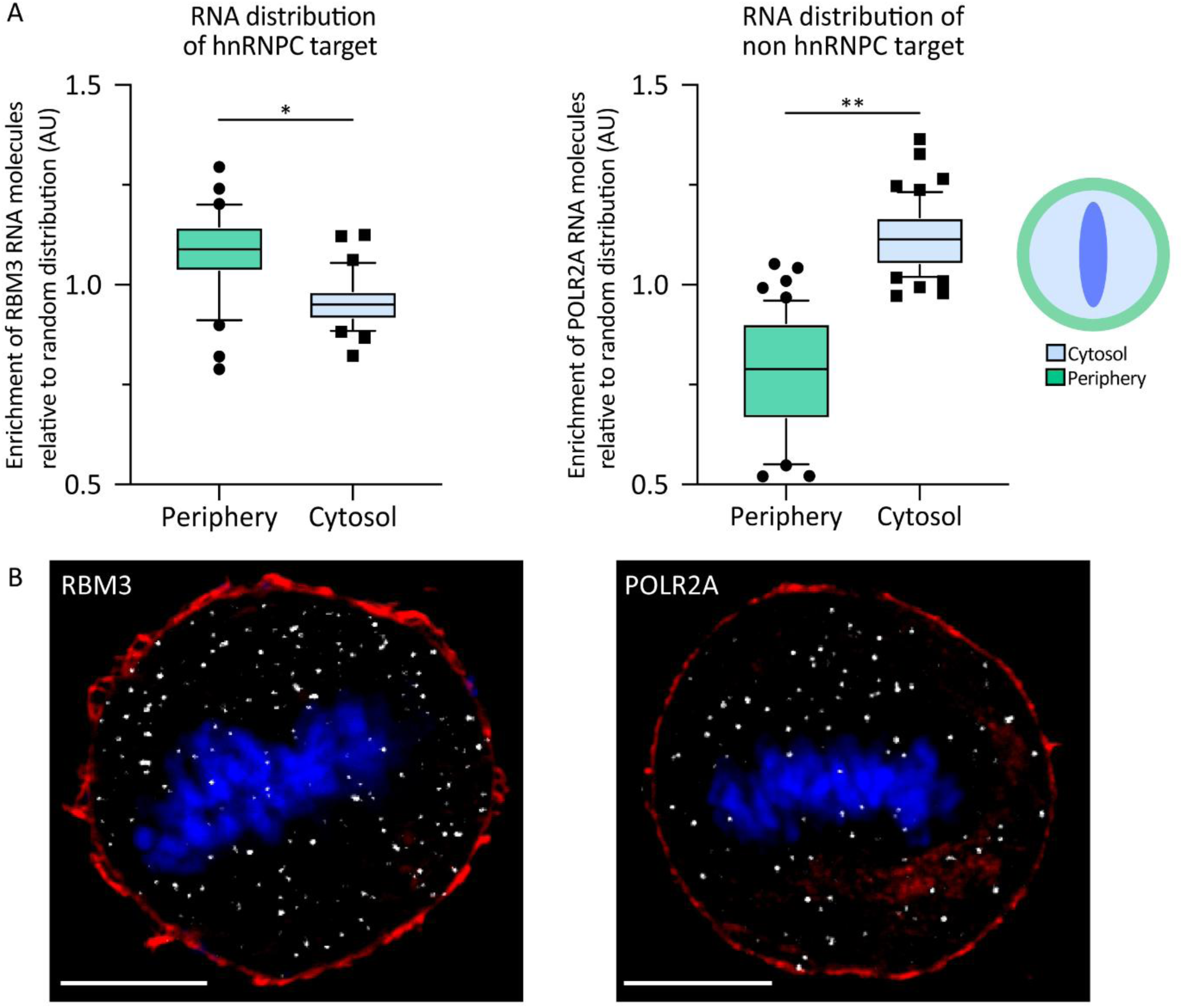
RBM3 and POLR2A mRNA subcellular distribution by smFISH. (A) smFISH signal enrichment in the cell periphery and in the remaining cytoplasm of HeLa S3 cells synchronized to metaphase (see definitions in Materials and Methods). Box and whiskers, 10-90 percentiles, of 3 independent experiments are shown. N_RBM3_=70, N_POLR2A_=112. Paired t-test: *P = 0.0002, **P < 0.0001. (B) Representative confocal smFISH images stained for single RNA molecules of either RBM3 or POLR2A (smFISH probes, white), actin (phalloidin, red), DNA (Hoechst, blue). Scale bar, 10 μm.

In a previous attempt to probe for mitotic hnRNPC’s role in post-transcriptional gene regulation, we generated gene expression (RNA-seq) datasets in DTB-synchronized cells, in the presence of DOX-induced shRNA-mediated hnRNPC knockdown (KD). These conditions could only achieve ∼50% reduction in hnRNPC levels, and similarly, cell synchronization was effective for ∼50% of cells (24); however still provided valuable insight. Here, we crossed mitotic differential mRNA expression in response to hnRNPC KD with hnRNPC targets (this study), to assess whether hnRNPC might affect the cellular levels of its specific mRNA targets. Evidently, even with only ∼50% hnRNPC depletion, the cellular levels of its targets were compromised, with no effect on levels of non-hnRNPC targets (Figure 6A). Specifically, upon hnRNPC KD, ∼60% of LD-hnRNPC and ∼70% of HD-hnRNPC exonic targets exhibited reduced steady-state RNA levels (p<0.0001, Figure 6A). Similarly, 65% of LD-hnRNPC intronic targets also reduced their steady state RNA levels in response to hnRNPC KD (p<0.0001, Figure 6B, left panel). This significant decline in levels of mitotic hnRNPC targets underscores an unexpected, yet critical dependency of transcriptome maintenance on hnRNPC during mitosis. To also probe whether hnRNPC affects the association of its mitotic targets with ribosomes, we crossed mitotic hnRNPC targets with ribosome footprinting (Ribo-seq) data in response to hnRNPC KD (24). Our analysis indicated a lack of major effect of mitotic hnRNPC expression level on the number of ribosome-protected fragments (RPF) along the exonic and intronic targets of hnRNPC, with a similar trend observed for non-hnRNPC targets (Figure 6AB). This observation hints at a more complex view (discussed below) stemming from the multistep nature of mitosis, where hnRNPC nuclear egress may either precede or follow the onset of RNA metabolism arrest, as well as the loading of recently matured mRNA with the translation machinery, which involves ribosome protection.

**Figure 6:**
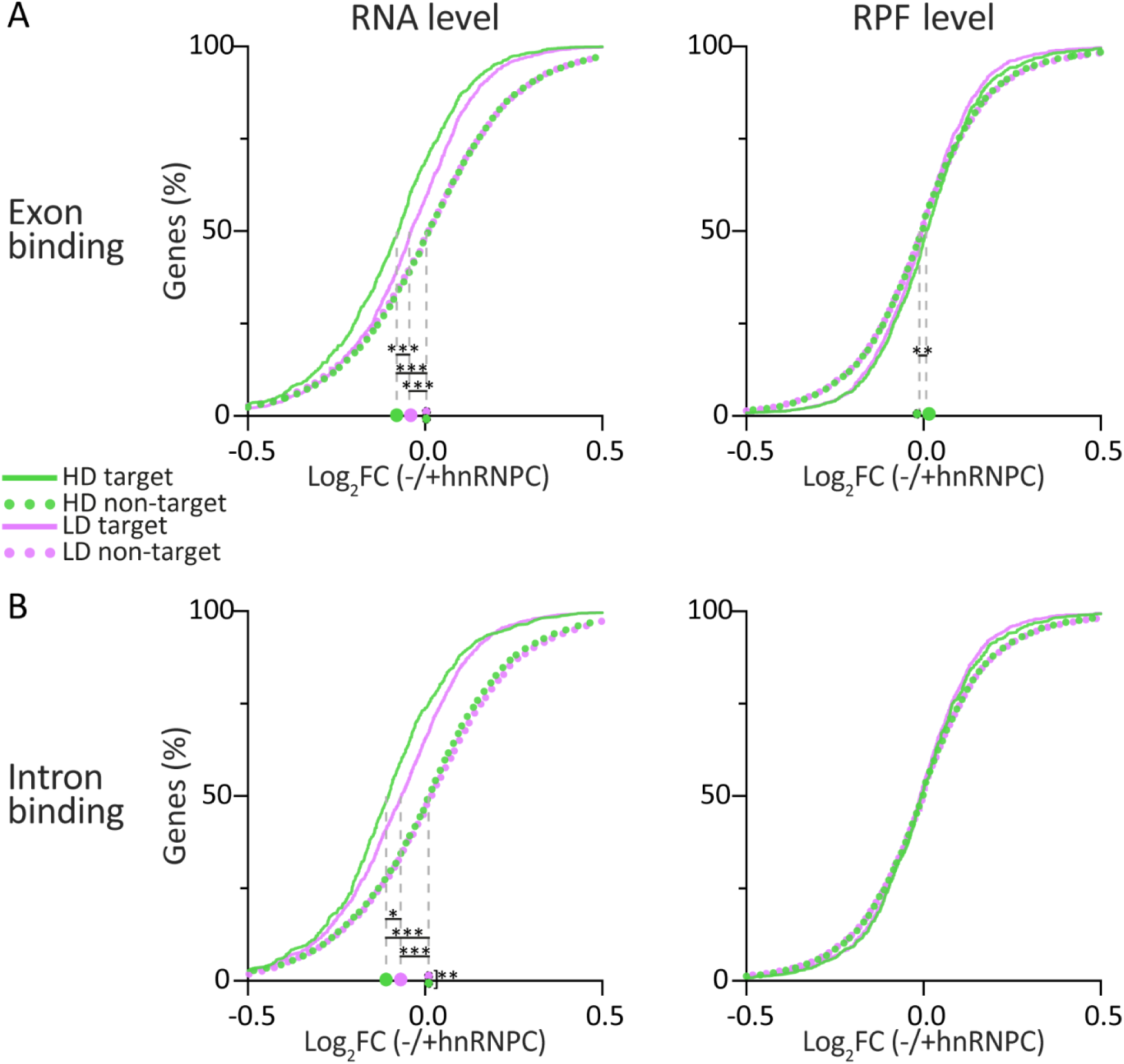
Effect of hnRNPC KD on mRNA levels and ribosome occupancy of hnRNPC targets and non-targets. Cumulative distribution of the change in RNA level (left), and ribosome-protected fragments (RPF, right) following DOX-mediated hnRNPC downregulation in cells synchronized to mitosis by DTB {Aviner, 2017 #256}. Shown is log_2_FC(-/+hnRNPC) of exonic (A) and intronic (B) hnRNPC targets and non-targets (this study). HD, green; LD, pink; Targets, continuous line; non-targets, dashed line. Median values are presented on the x-axis. Kruskal–Wallis test: *P = 0.02, **P = 0.01, ***P < 0.0001.

Collectively, the data point to the global role of hnRNPC as a mitotic pre-mRNA and mRNA stabilizer. To search for additional insights related to specific genes, we focused on a list of 229 genes expressed at sufficient levels in our different datasets, enabling reliable comparisons (Supplementary Figure S1C and Supplementary Table S1). This list includes 52 genes that embrace LD and/or HD exonic and/or intronic hnRNPC binding sites. We noticed that among the genes showing ≥20% reduction in RNA abundance upon hnRNPC KD, the decrease in RPF values does not always follow similar trends. For example, the RNA and RPF values of TAF1D decreased by ∼25% and ∼15%, respectively. However, the effect of hnRNPC KD on association with ribosomes revealed an opposite direction for SLC30A1, which exhibits a reduction of ∼23% in RNA level and an increase of ∼12% in RPF. While the latter is a rarer case, it points to an interesting possibility, which is discussed below.

## Discussion

Precise cell cycle regulation requires an accurate gene expression via multiple regulatory mechanisms, which include RNA stability. Upon entry to mitosis, RNA metabolism is broadly suppressed by a significant halt of transcription, splicing, RNA degradation, and translation (1–10). This suppressed RNA metabolism likely enables the cell to redirect essential resources towards proper chromatin separation, the central event in mitosis. The scheduled degradation of specific pre-mRNA/mRNA populations at the mitosis-to-G1 phase transition (63) raises the question of how the rest of the transcriptome is protected upon nuclear envelope breakdown. RBPs evolved to interact with their specific RNA recognition elements (RREs); however, despite the specificity at the molecular level, there are examples of the multifunctionality of a given binding specificity in regulating distinct biological functions along the mRNA life and at different cellular contexts (14–23). Here, we used an unbiased approach to specifically focus on hnRNPC, an abundant nuclear RBP that interacts mainly with introns within the nucleus and is largely involved in splicing during interphase. An iCLIP experiment using non-synchronized cells revealed that only ∼10% of hnRNPC’s binding targets are within exons (27). Our results, obtained using cells synchronized to metaphase, show that between 36-56% of hnRNPC’s binding sites are within exons (Figure 3B). We found that while mitotic hnRNPC continues to interact with its established U-rich RRE (27,53–55), it acquires distinct targets, in addition to those previously reported.

Known as a regulator of alternative splicing during interphase, hnRNPC was also implicated in the sporadic cases of 3’UTR binding and stabilization of urokinase receptor (ukR) and amyloid precursor protein (APP) mRNAs, thereby enhancing their translation (64,65). Here, we exposed a global role of mitotic hnRNPC as a general RNA stabilizer. We show that downregulation of hnRNPC elicited a negative effect on the abundance of 60%-70% of its mitotic targets (Figure 6). This shared effect of interactions with both pre-mRNAs and mature mRNAs, indifferent to the distinct complexes harboring these transcripts, aligns with the necessity to keep the entire mitotic transcriptome stable upon the inhibition of RNA metabolism and loss of compartmentalization at M-phase, before the degradation of specific RNA molecules towards mitosis exit occurs (63). Due to the limitation of cell synchronization, we could not show RNA stabilization directly by introducing transcription inhibitors such as Actinomycin D (66). However, the natural suppression of transcription during mitosis (5) may effectively replace transcription inhibition, rendering steady-state mRNA generally representing its stability.

Using sucrose gradient fractionation of metaphase-synchronized cells, we separately characterized HD-hnRNPC, which exhibited predominant interaction of hnRNPC with the 3’UTR of spliced mature mRNAs, in the context of poly-ribosomes. In addition, we characterized LD-hnRNPC, which co-sedimented with mono-ribosomes and spliceosomes. Our analyses indicate that LD-hnRNPC interactions represent the blended nature of this fraction. Specifically, exonic targets of LD-hnRNPC were predominantly mature mRNAs with shorter CDS lengths, as would match loading with single ribosomes. The LD-hnRNPC intronic targets co-sediment with spliceosomes, as would match hnRNPC-bound pre-mRNAs. Importantly, we confirmed that the binding sites within all analyzed subsets of hnRNPC targets display their established U-rich RRE (Figure 3A; Supplementary Figure S3B). Three key findings strongly support the authenticity of our conclusion that mitotic hnRNPC acquires interactions with mature mRNAs, in addition to unprocessed pre-mRNAs. These include: (i) the length dependence feature of hnRNPC exonic interactions is in accord with its reported cooperative binding property (Figure 4B); (ii) the 3’UTR being the predominant binding site of hnRNPC exonic targets (Figure 3B, 4E) is in accord with authentic interactions with mature mRNAs engaged with ribosomes along their CDS; (iii) the finding that it is the 3’UTR length that limits hnRNPC cooperative binding (Figure 4C, D).

To obtain our observations, we were highly invested in optimal cell synchronization. Our efforts were designed to obtain preparative amounts of cells, at highly homogenous synchronization to the metaphase step. These enabled fCLIP analysis in conditions that minimize interphase-contamination and sub-M-phase diversity. While our results may reflect a snapshot representing a single time point in mitosis, they revealed a dramatic functional shift in detected hnRNPC interactions. Such a functional change might be shared by other RBPs, including highly abundant nuclear RBPs during mitosis. Such dynamic remodeling of protein assemblies on RNA in response to physiological signals in living cells, thereby affecting the function of RBPs was recently demonstrated (67), though not in a cell cycle-specific context.

hnRNPC exists as two isoforms, hnRNPC1 and hnRNPC2, produced through alternative splicing. Both isoforms form a stable tetramer in vivo, predominantly in a (C1)_3_ (C2)_1_ ratio (68). Importantly, the 13-amino-acid insertion in hnRNPC2 influences RNA-binding affinity and specificity (69). The antibody used in this study does not discriminate between both isoforms, thus allowing us to obtain an integrated, physiologically representative profile that likely reflects the natural hnRNPC binding landscape within the mitotic cell. Post-translational modifications of hnRNPC isoforms, such as phosphorylation and sumoylation, affect their binding capability to RNA and other protein partners (64,70) and may also affect tetrameric composition, which remains to be studied in the context of mitotic hnRNPC function.

Our early work has shown that despite the significant reduction in active translation in cells synchronized to mitosis using DTB, they are generally immune to stress granule formation. In addition, we found that despite the decrease in translation, DTB-synchronized cells harbor heavy poly-ribosomes, of which some are actively translating, whereas others are attenuated/stalled (6). Although the mechanism governing transcript-specific active or attenuated elongation is unknown, the findings imply that ribosome association with mitotic transcripts leads to their protection. The current study informs that hnRNPC’s downregulation negatively affects the abundance of its mitotic targets but does not significantly impact their RPF value (Figure 6AB). This observation implies that the RNA protection property of mitotic hnRNPC is more significant for the transcripts that are not fully engaged with ribosomes. The fCLIP data allows the estimation of each mitotic hnRNPC target’s status in metaphase. As observed in Supplementary Table S3, upon hnRNPC KD, most cases exhibiting a decrease in RNA abundance demonstrate either no change or minimal reduction in RPF value. In addition to its global function as an RNA stabilizer, looking into individual targets (IGV maps are presented in Supplementary Figure S6) suggests that mitotic hnRNPC may also exert gene-specific regulatory effects. For example, upon hnRNPC KD, the TAF1D transcript shows a decrease in RNA abundance and RPF by 25% and 15%, respectively (Supplementary Table S3). This could reflect the greater negative impact of hnRNPC KD on the stability of the TAF1D RNA sub-population that is not sufficiently protected by ribosomes. On the other hand, the fact that reduction of hnRNPC abundance also leads to some decrease in ribosome association could hint at the possibility that hnRNPC has a positive regulatory role on the translation of specific transcripts during mitosis, as previously shown for c-Myc and Unr mRNAs, among others (24,71,72). Interestingly, TAF1D encodes a subunit of RNA polymerase I. It is vital for rRNA transcription, ribosome biogenesis, protein synthesis, and cell cycle progression. Moreover, its expression is maintained constant throughout the cell cycle and is subjected to cell cycle-dependent translational control (73). Importantly, its downregulation leads to downregulation of genes important for mitosis, such as CDK1 (74). Therefore, collectively, it is tempting to speculate that hnRNPC contributes to the active translation of TAF1D mRNA in mitotic cells. A possible mechanism of promoting translation might be by relieving the translational block elicited by other inhibitory RBPs via their displacement. The fCLIP data also points to more rare cases that exhibit a decrease in RNA abundance and an increase in the RPF value. For example, upon hnRNPC KD, the SLC30A1 transcript shows a decrease in RNA abundance by ∼23% and an increase in RPF by 12% (Supplementary Table S3). The latter suggests an increased binding of ribosomes upon hnRNPC downregulation, implying a possible translation-rate compensatory mechanism for the reduced levels of mRNA, or an inhibitory role for hnRNPC on active SLC30A1 mRNA translation during mitosis. The SLC30A1 gene encodes Zinc Transporter 1, predominantly localized to the plasma membrane. Zinc regulates various enzymes and calcium-dependent cellular processes and is important for cell cycle control (75). According to our data, it is tempting to speculate that hnRNPC is involved in the translation inhibition of SLC30A1 mRNA in mitotic cells. A possible mechanism of inhibiting translation might be by blocking ribosome association. Together, we suggest the possibility that, in addition to the function of hnRNPC as an RNA stabilizer during mitosis, it may also acquire additional functions as a translation regulator, provided by its various protein partners. Future experiments will identify the different hnRNPC-RNPs partners, their specific post-translational modifications, the specific signals that govern their dynamic changes, and decipher the functional consequences.

Importantly, hnRNPC dysregulated activity correlates with poor prognosis in multiple malignancies (76). Our results expand the known functions of hnRNPC, proposing a model in which, aside from splicing, mitotic hnRNPC promotes global transcriptome maintenance during mitotic suppression of RNA metabolism, which can explain its specific contribution to rapidly dividing cancer cells. Evolvement of preservative mechanisms to maintain transcriptome integrity during this sensitive phase in the cell cycle to provide the daughter cells with an economical and efficient start as they enter G1 phase may represent a broader manifestation and vulnerability of cancer cells. Future studies should be designed to provide additional insights into hnRNPC’s multifunctional roles during mitosis.

## Acknowledgement

We thank Markus Hafner for fruitful discussions.

## Supplementary Figures

**Figure S1:**
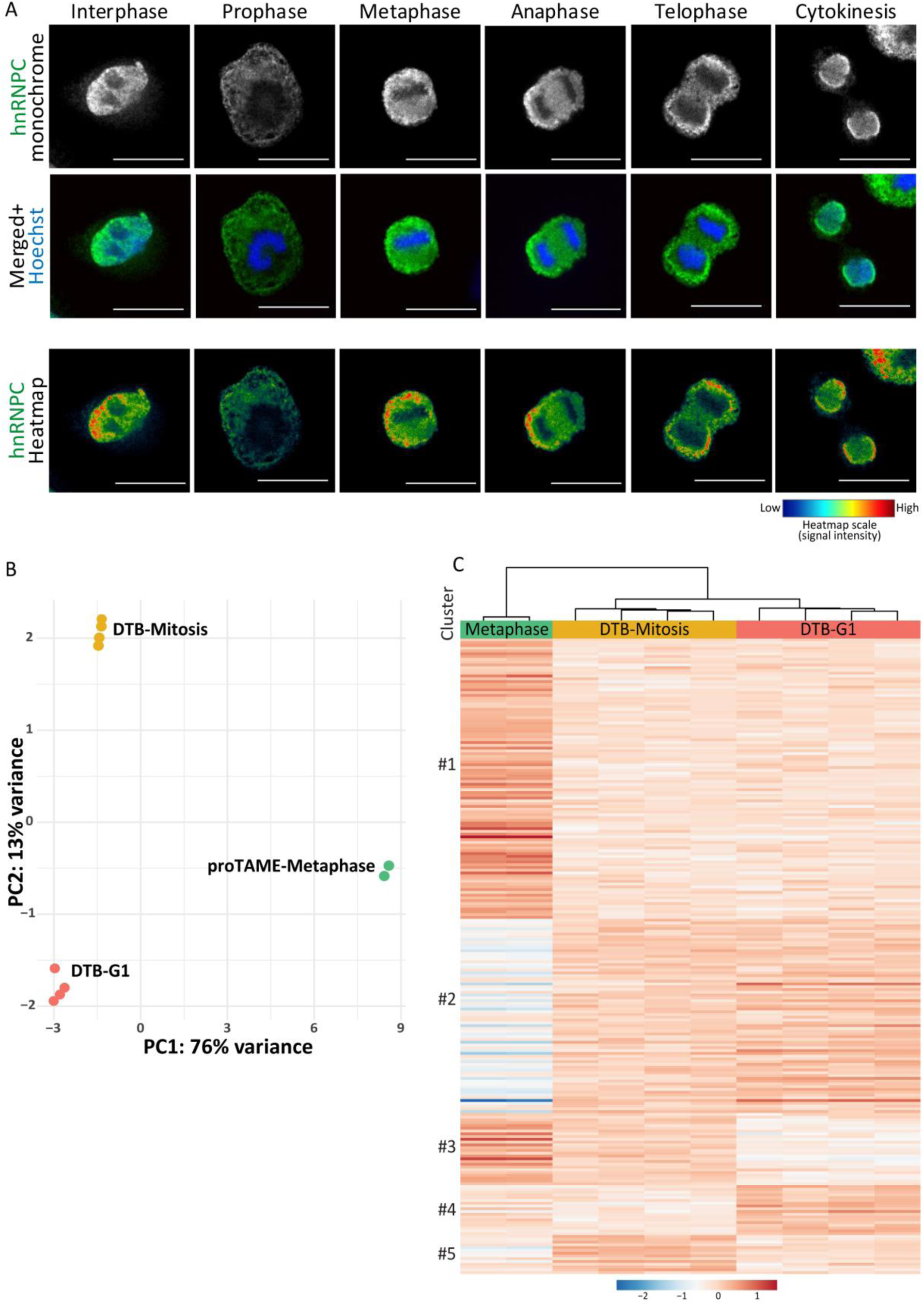
Cell cycle and gene expression analyses. (A) Representative images of hnRNPC subcellular localization in interphase and during different Mitosis substages. Top: hnRNPC (green), Middle: hnRNPC merged with DNA (Hoechst, blue), Bottom: Pseudocolor heatmap of hnRNPC signal. Heatmap color scale attached. Scale bar, 40 μm. (B) Principal Component Analysis (PCA) of variance stabilizing transformation (VST) values of 229 filtered genes in cells synchronized by ProTAME to Metaphase (current study), and by DTB to mitosis or G1 *{Aviner, 2017 #256}* (See standardization and comparison to RNAseq data in Materials and Methods and supplementary Table 1). Each point represents a biological replicate. (C) Heatmap of unsupervised hierarchically clustered row-centered VST gene expression data (as in (B)).

**Figure S2:**
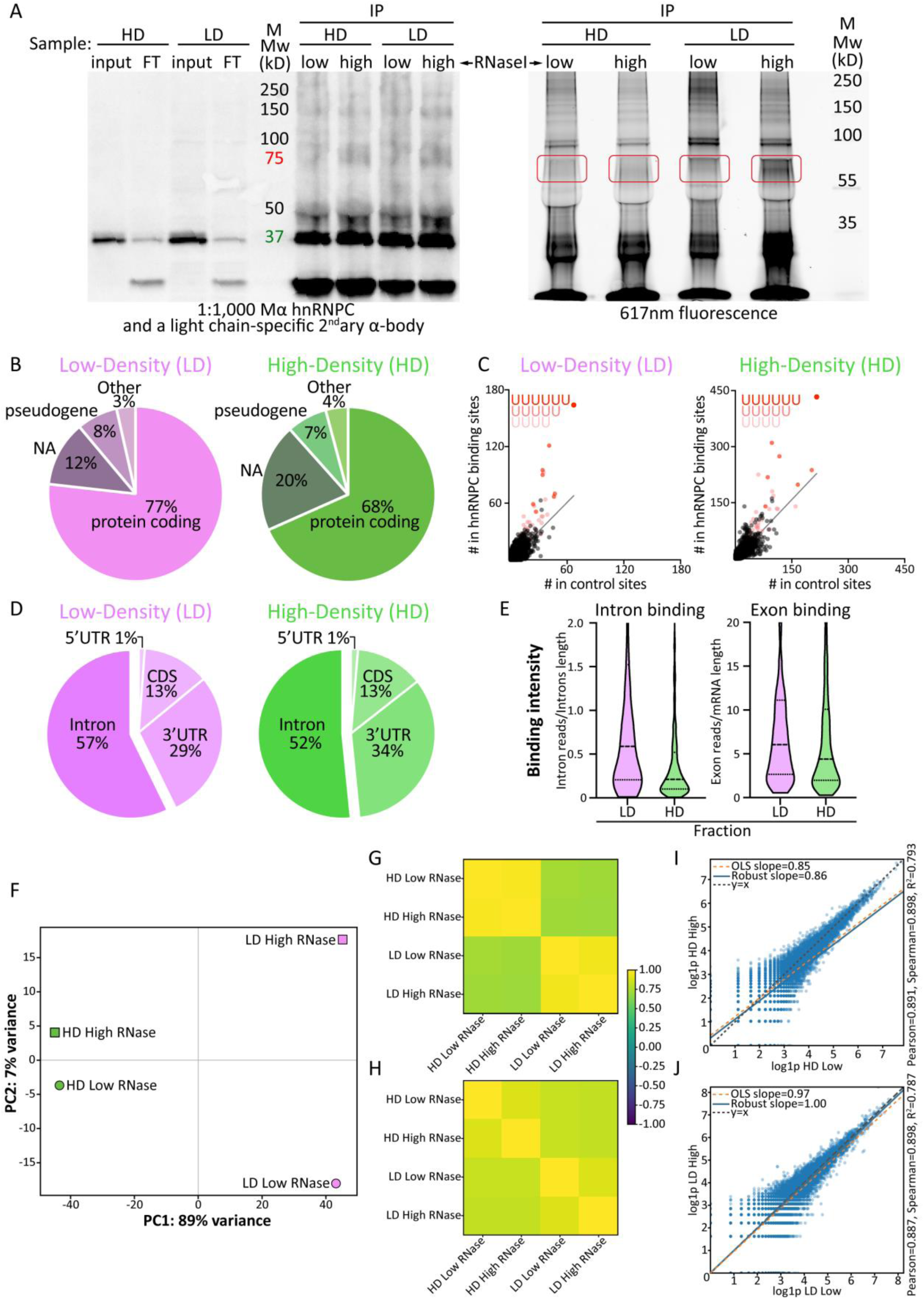
fCLIP procedure and replicates analysis. (A) Immunoblots (left) and 617nm fluorescence imaging (right) of hnRNPC immunoprecipitation from low-density (LD) and high-density (HD) sucrose gradient fractions, treated with 0.015 U/μL (low) or 0.15 U/μL (high) RNase I. Input, flowthrough (FT), and immunoprecipitated (IP) material are shown. Red boxes mark regions excised for downstream library preparation. (B) Distribution of hnRNPC binding sites (groups) by transcript annotation in LD and HD fractions (NA-no annotation). Refers to the low-RNase treatment <u>(C-E) refer to the high RNase treatment as follows:</u> (C) 6-mer analysis for sequence motif enrichment within hnRNPC binding sites. Scatterplots show observed frequencies of each 6-mer relative to its frequency in a control set. 6-mers containing 4, 5, and 6 consecutive U are color labelled. (D) Distribution of hnRNPC binding between introns and exons, segmented by mRNA functional elements. (E) hnRNPC binding intensity per intronic and exonic binding sites, calculated per gene by normalizing read counts to introns or exons length <u>(F-J) refers to low- and high-RNase data (total of 4 datasets) as follows:</u> (F) PCA of fCLIP replicates, filtered for protein coding transcripts. (G,H) Inter-sample Pearson (G) and Spearman (H) correlation heatmaps, across replicates (LD-hnRNPC - Pearson=0.96; Spearman=0.90, and HD-hnRNPC - Pearson=0.98; Spearman=0.90). (I,J) Scatterplots of log1p-transformed read counts (*statsmodels*) for HD (I) and LD (J) replicates. Ordinary least squares (OLS) and robust regressions are plotted.

**Figure S3:**
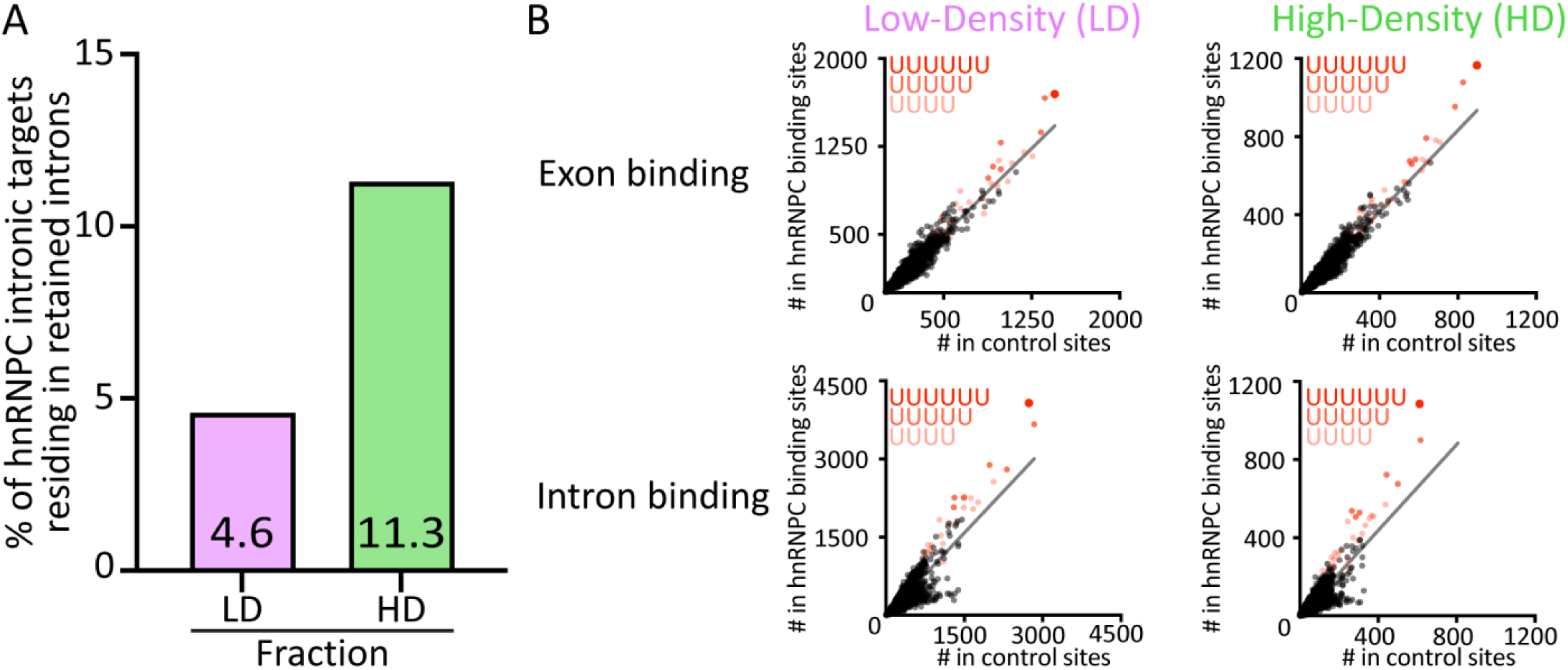
Characterization of hnRNPC binding sites. (A) Proportion of hnRNPC intronic binding sites residing in previously reported retained introns {Mercer, 2015 #77}. (B) 6-mer analysis for sequence motif enrichment within LD- and HD-hnRNPC-bound exonic and intronic regions. Scatterplots show observed frequencies of each 6-mer relative to its frequency in a control set. 6-mers containing 4, 5, and 6 consecutive U are color labelled.

**Figure S4:**
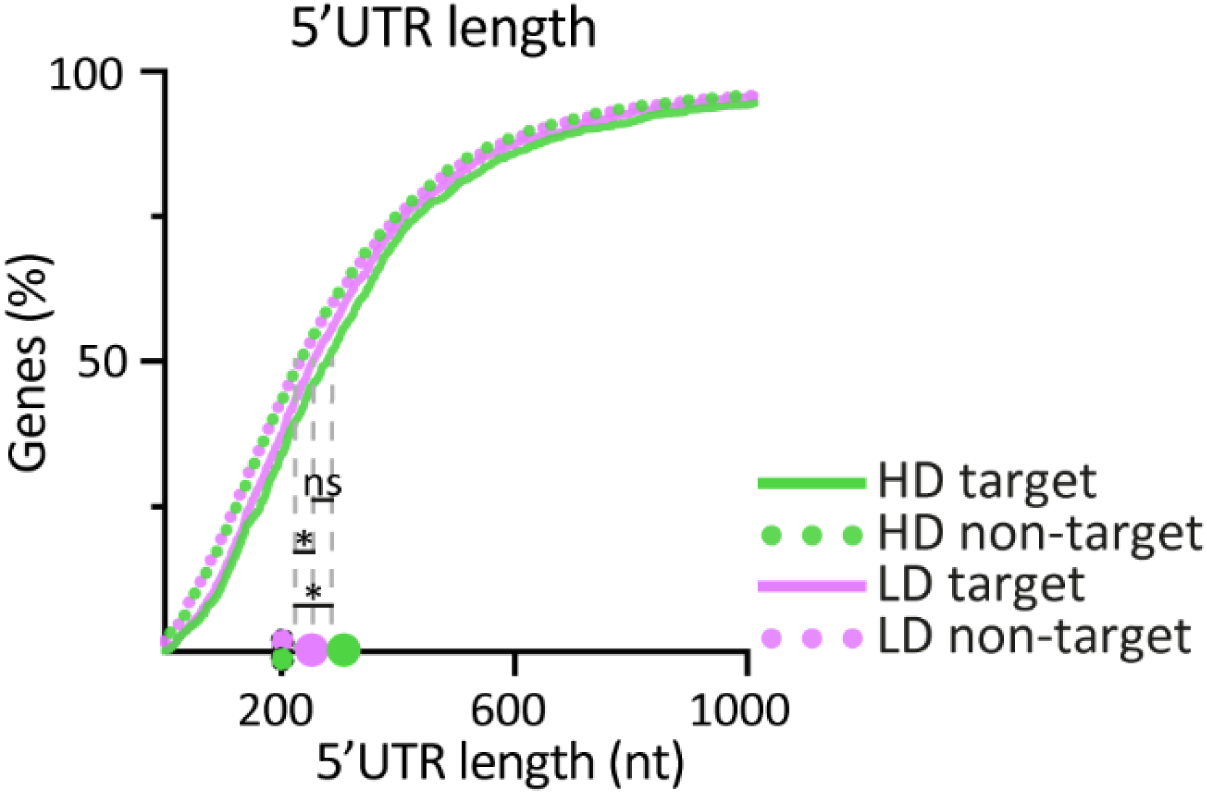
Analysis of hnRNPC length dependent Exonic Binding to 5’UTR. Cumulative 5’UTR length distribution of LD- and HD- hnRNPC targets and non-targets. P-values (Kruskal–Wallis): *P < 0.0001; ns-not significant, UTR-untranslated region

**Figure S5:**
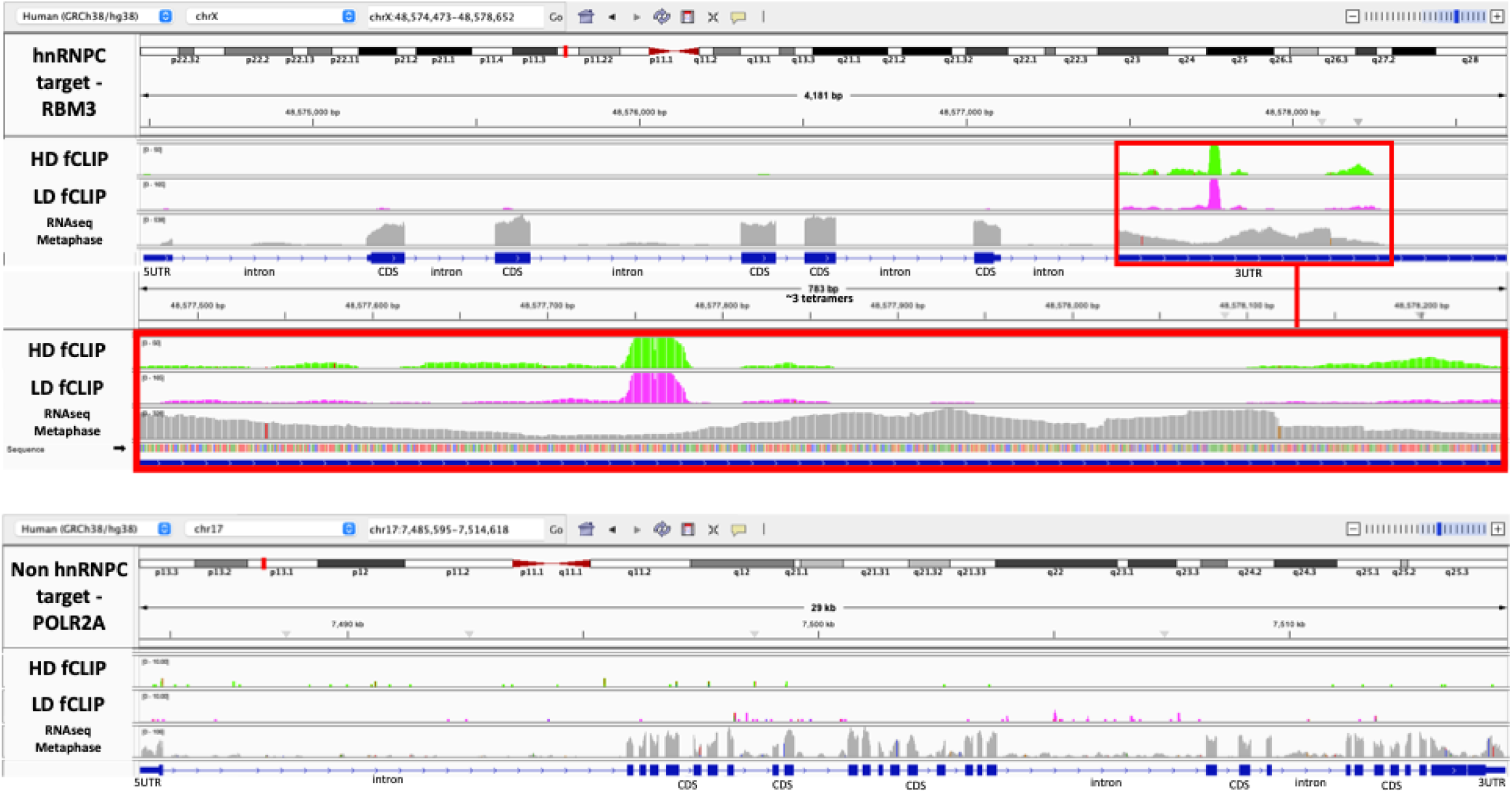
IGV visualization of hnRNPC-bound and non-bound transcripts used for smFISH experiments. Integrative Genomics Viewer (IGV) {Thorvaldsdottir, 2013 #101} screenshots showing hnRNPC fCLIP and RNA-seq raw reads along *RBM3* coordinates (top, hnRNPC target) and *POLR2A* coordinates (bottom, non-hnRNPC target). Coverage tracks are color-coded: Green, high-density (HD) hnRNPC fCLIP; Pink, low-density (LD) hnRNPC fCLIP; Grey, Metaphase cells total extract RNA-seq reads. In *RBM3*, a ∼783 nt binding region corresponding to the occupancy of ∼3 hnRNPC tetramers is highlighted (red box). Thin horizontal lines represent introns, filled boxes represent exons, with thicker boxes denoting the CDS and thinner boxes denoting UTRs. Gene orientation is indicated by the direction of the arrows along the connecting lines. CDS-coding sequence, UTR-untranslated region.

**Figure S6.**
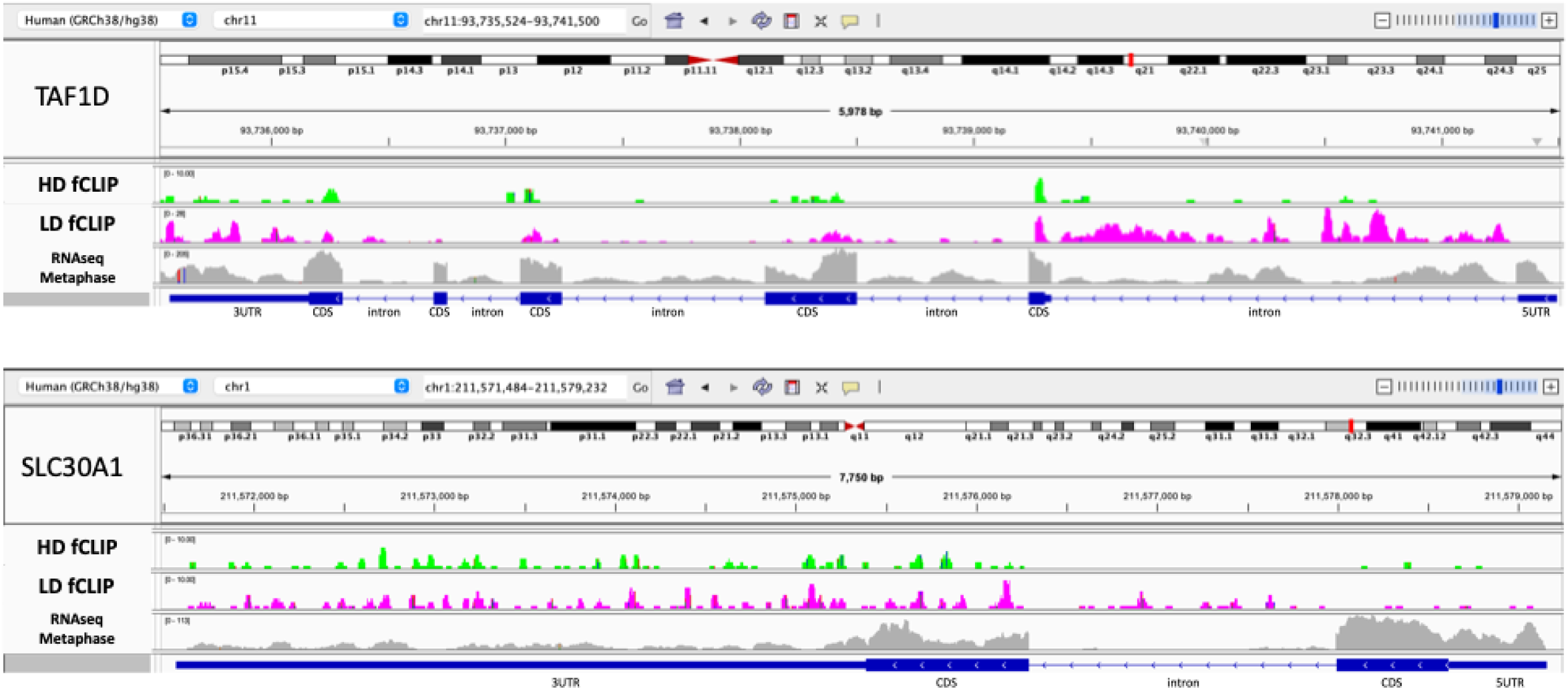
Integrative Genomics Viewer (IGV) {Thorvaldsdottir, 2013 #101} screenshots showing hnRNPC fCLIP and RNA-seq raw reads along *TAF1D* coordinates (top) and *SLC30A1* coordinates (bottom). Coverage tracks are color-coded: Green, high-density (HD) hnRNPC fCLIP; Pink, low-density (LD) hnRNPC fCLIP; Grey, Metaphase cells total extract RNA-seq reads. Thin horizontal lines represent introns, filled boxes represent exons, with thicker boxes denoting the CDS and thinner boxes denoting UTRs. Gene orientation is indicated by the direction of the arrows along the connecting lines. CDS-coding sequence, UTR-untranslated region.

